## Supplemental Material for "Network-wide abnormalities explain memory variability in hippocampal amnesia"

### Supplementary Material

#### S1: Composite Memory scores for LE patients and healthy controls

| **Memory Composite Score** | **Controls** | | **Patients** | | **Controls vs. Patients** | | |
| --- | --- | --- | --- | --- | --- | --- | --- |
|  | **Mean** | **SD** | **Mean** | **SD** | **Test** | **Statistic** | **p-corr** |
| Anterograde Retrieval (z) | 0.72 | 0.55 | -0.86 | 0.89 | Wt | 9.40 | < 0.0005 |
| Anterograde Retention (Visual Forgetting) (z) | 0.23 * | 0.36 | -0.52 * | 1.17 | U | 433.00 | 0.0005 |
| Remote Autobiographical (max = 18) | 15.93 * | 2.63 | 9.77 | 4.82 | U | 130.00 | < 0.0005 |

Table S1.1: Composite memory scores for healthy controls and LE patients; ‘max = 18’: the maximum score sum attainable for the autobiographical memory for childhood and early adulthood in the AMI (Kopelman et al., 1989), which does not use age-scaled / standardized scores; p-corr: p values are adjusted for multiple testing using the Holm-Bonferroni sequential method (n=3); U: Mann-Whitney U; Wt: Welch’s test; z: standardized age-scaled scores; *: Shapiro-Wilk test: p < 0.05.

#### S2: Relationship of HPC atrophy with PCC functional abnormalities

Given the reciprocal connectivity of thalamic nuclei with both the HPC and the cingulate cortex (Aggleton, 2014; Aggleton et al., 2010; Bubb et al., 2017) and our hypothesis that HPC atrophy is followed by structural and functional abnormalities in interconnected areas within the HPC-diencephalic-cingulate networks, we assumed a causal chain of events, whereby HPC damage has remote effects on thalamic volume, which, in turn, leads to abnormalities in the cingulate cortex, observed here in the form of reduced rsALFF in the PCC. We used a series of bivariate correlations and mediation analyses, with HPC volume as the independent variable, thalamic volume as the mediator variable, and rsALFF in the PCC as the dependent variable. A mediation analysis supported this hypothesis, showing that the effects of the average GM volume reduction in the HPC VBM clusters on patients’ reduced rsALFF in the PCC were fully mediated by the correlative reduction of the average GM volume of the thalamic VBM clusters (direct effect: β=0.26, p=0.064; indirect effect: β=0.16, 95% CI: 0.004,0.403; figure S2.1).

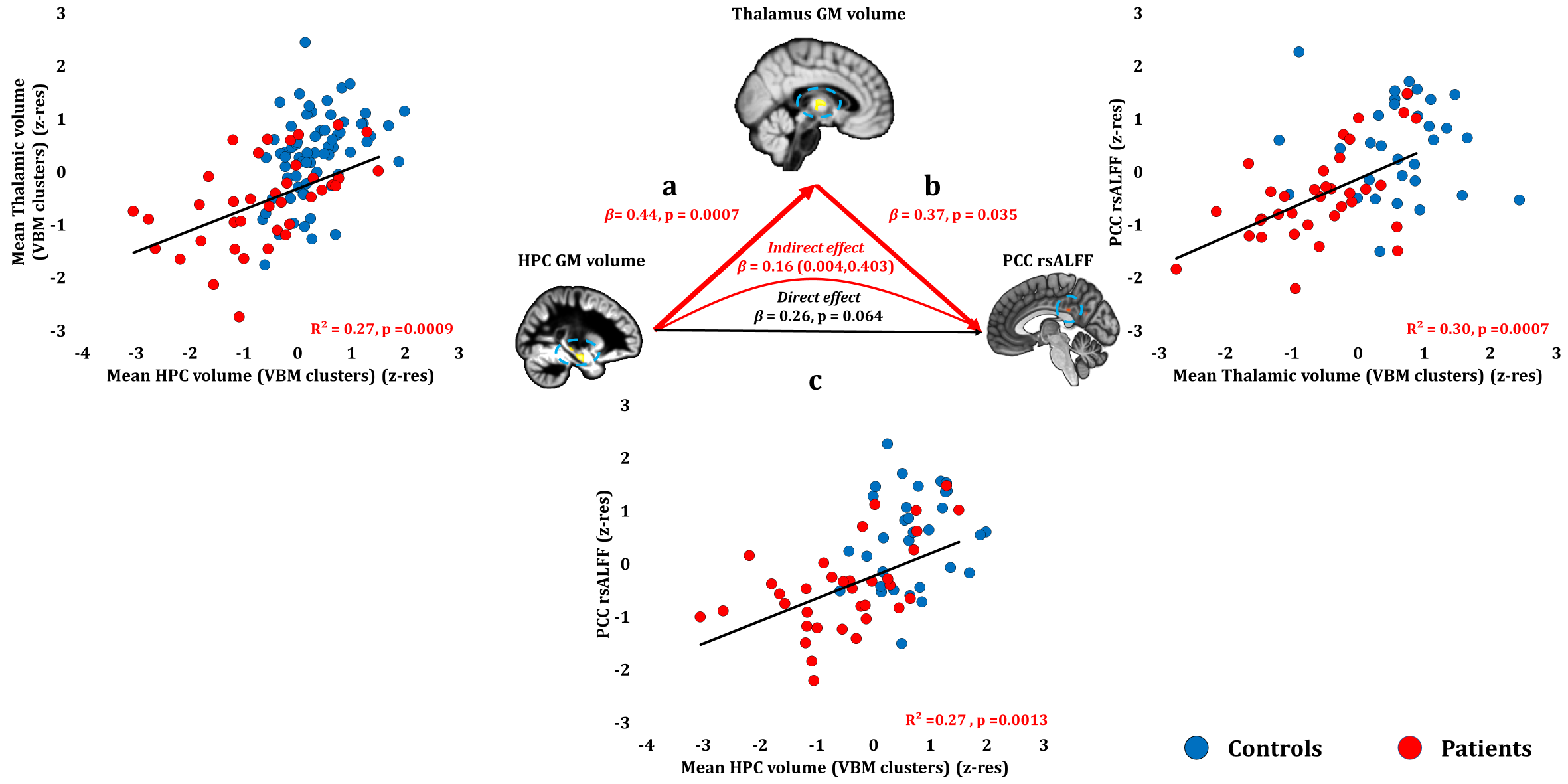

Figure S2.1: Relationship of HPC atrophy with PCC rsALFF reduction across patients: mediation analysis: a: mean GM volume of the two HPC clusters correlated with the mean GM volume of the two thalamic clusters across patients; b: mean PCC rsALFF correlated across patients with the mean GM volume of the two thalamic clusters; the mediation analysis demonstrates that this effect held when the correlation of thalamic GM volume with the mean GM volume of the HPC clusters was accounted for; c: mean GM volume of the HPC clusters correlated with PCC rsALFF; the mediation analysis demonstrated that this relationship did not hold over and above the correlation of the mean GM volume of the thalamic clusters with the HPC clusters; there was thus no direct effect of reduced HPC GM volume on PCC rsALFF (within parenthesis: 95% confidence intervals); GM: grey matter; HPC: hippocampus; MAP: Memory and Amnesia Project; OPTIMA: Oxford Project To Investigate Memory and Aging; rsALFF: resting-state amplitude of low frequency fluctuations; TIV: total intracranial volume; VBM: voxel-based morphometry; z-res: GM volumes from VBM clusters are residualized against age, sex, scan source (MAP, OPTIMA), and TIV across participants; mean rsALFF is residualized against age and sex across participants.

#### S3: Relationship of delay since symptom onset with structural and functional abnormalities

Given our hypothesis that broader network abnormalities unfold as a consequence of acute HPC atrophy (i.e. at a time-point after the acute focus of HPC damage), we also assessed the relationship of the delay between symptom onset and research participation with the extent of HPC and extra-HPC structural and functional abnormalities in a series of bivariate correlations. Of the structural/functional abnormalities (n=13) identified in our patient group, only inter-HPC rsFC decreased across patients as a function of the delay between symptom onset and research scan (rho=-0.58, p-corr=0.004). No other brain abnormalities showed this relationship, even at uncorrected levels (all rhos, |rho| ≤ 0.255; all ps, p-unc ≥ 0.122; table S3.1).

| **Measure (z-res)** | **rho** | **p-corr** |
| --- | --- | --- |
| **Inter-HPC rsFC** | **-0.578** | **0.0036** |
| R HPC volume (VBM) | -0.255 | > 0.9999 |
| L HPC volume (VBM) | -0.216 | > 0.9999 |
| R HPC volume (segmentation) | -0.202 | > 0.9999 |
| PrCu rsALFF | -0.120 | > 0.9999 |
| PCC rsALFF | 0.098 | > 0.9999 |
| L HPC volume (segmentation) | -0.089 | > 0.9999 |
| Medial thalamus (VBM) | -0.081 | > 0.9999 |
| R HPC - PCC rsFC | -0.082 | > 0.9999 |
| R thalamus volume (VBM) | -0.038 | > 0.9999 |
| R ERC volume (segmentation) | -0.013 | > 0.9999 |
| R HPC-MPFC rsFC | 0.009 | > 0.9999 |
| L thalamus volume (segmentation) | -0.003 | > 0.9999 |

Table S3.1: Bivariate correlations of the delay between symptom onset and research participation with volumes, mean rsALFF and mean rsFC of clusters where patients showed structural and functional abnormalities; **bold**: p-corr < 0.05; HPC: hippocampus; L, R: left, right (hemisphere); MPFC: medial prefrontal cortex; p-corr: p values are adjusted for multiple testing (n=13) using the Holm-Bonferroni sequential method; PCC: posterior cingulate cortex; PrCu: precuneus; rho: Spearmann’s rank correlation coefficient; rsALFF: resting-state amplitude of low frequency fluctuations; rsFC: resting-state functional connectivity; OPTIMA: Oxford Project To Investigate Memory and Aging; Memory and Amnesia Project; TIV: total intracranial volume; z-res: volumes are residualized against age, sex, scan source (MAP, OPTIMA), and TIV; functional abnormalities are residualized against age and sex;

#### S4: Correlation of memory scores with structural and functional abnormalities across patients

| **Composite Score (z)** | **Structural/Functional abnormalities (z-res)** | **r/rho** | **p-corr** |
| --- | --- | --- | --- |
| **Anterograde Retrieval (z)** | L HPC volume (segmentation) | 0.274 | > 0.999 |
|  | L HPC volume (VBM cluster) | 0.462 | 0.105 |
|  | L thalamus (segmentation) | 0.241 | > 0.999 |
|  | M thalamus volume (VBM cluster) | 0.361 | 0.598 |
|  | PCC rsALFF | 0.551 | 0.024 |
|  | PrCu rsALFF | 0.354 | 0.684 |
|  | R ERC volume (segmentation) | 0.271 | > 0.999 |
|  | R HPC - MPFC rsFC | 0.051 | > 0.999 |
|  | R HPC - PCC rsFC | 0.281 | > 0.999 |
|  | R HPC volume (segmentation) | 0.290 | > 0.999 |
|  | R HPC volume (VBM cluster) | 0.342 | 0.684 |
|  | R thalamus volume (VBM cluster) | 0.376 | 0.540 |
|  | R-L HPC rsFC | 0.389 | 0.546 |
| **Anterograde Retention (Visual Forgetting) (z)** | L HPC volume (segmentation) | 0.412 | 0.466 |
|  | L HPC volume (VBM cluster) | 0.401 | 0.532 |
|  | L thalamus volume (segmentation) | 0.288 | > 0.999 |
|  | M thalamus volume (VBM cluster) | 0.179 | > 0.999 |
|  | PCC rsALFF | 0.399 | 0.576 |
|  | PrCu rsALFF | 0.281 | > 0.999 |
|  | R ERC volume (segmentation) | 0.365 | 0.680 |
|  | R HPC - MPFC rsFC | 0.290 | > 0.999 |
|  | R HPC - PCC rsFC | 0.055 | > 0.999 |
|  | R HPC volume (segmentation) | 0.508 | 0.079 |
|  | R HPC volume (VBM cluster) | 0.556 | 0.024 |
|  | R thalamus volume (VBM cluster) | 0.338 | 0.850 |
|  | R-L HPC rsFC | 0.122 | > 0.999 |
| **Remote Autobiographical Memory (18)** | L HPC volume (segmentation) | 0.393 | 0.608 |
|  | L HPC volume (VBM cluster) | 0.467 | 0.272 |
|  | L thalamus volume (segmentation) | 0.558 | 0.041 |
|  | M thalamus volume (VBM cluster) | 0.463 | 0.297 |
|  | PCC rsALFF | 0.448 | 0.493 |
|  | PrCu rsALFF | 0.460 | 0.434 |
|  | R ERC volume (segmentation) | 0.025 | > 0.999 |
|  | R HPC - MPFC rsFC | 0.227 | > 0.999 |
|  | R HPC - PCC rsFC | 0.291 | > 0.999 |
|  | R HPC volume (segmentation) | 0.395 | 0.608 |
|  | R HPC volume (VBM cluster) | 0.413 | 0.546 |
|  | R thalamus volume (VBM cluster) | 0.463 | 0.297 |
|  | R-L HPC rsFC | 0.107 | > 0.999 |

Table S4.1: analysis-level correction for multiple testing (n=39); **bold**: p-corr < 0.05; GM: grey matter; HPC: hippocampus; L: left hemisphere; M: medial; MAP: Memory and Amnesia Project; MPFC: medial prefrontal cortex; OPTIMA: Oxford Project To Investigate Memory and Aging; PCC: posterior cingulate cortex; p-corr: p values of bivariate correlations are corrected for multiple testing using the Holm-Bonferroni sequential method of correction for the total number of correlations (n = 39); PMC: posteromedial cortex; PrCu: precuneus; rsALFF: amplitude of low frequency fluctuations; rsFC: resting-state functional connectivity; R: right hemisphere; TIV: total intracranial volume; VBM: voxel-based morphometry; z-res: Mean rsALFF and rsFC values are residualized for age and sex across participants; volumes are residualized against age, sex, scan source (OPTIMA, MAP) and TIV across participants.

| Structural/Functional  abnormalities (z-res) | Verbal Recognition (z) | | Visual Recognition (z) | | Verbal Recall (z) | | Visual Recall (z) | | Visual Forgetting (z) | | Remote Autobiographical (18) | |
| --- | --- | --- | --- | --- | --- | --- | --- | --- | --- | --- | --- | --- |
|  | r | p-corr | r | p-corr | r | p-corr | Rho | p-corr | rho | p-corr | r | p-corr |
| PCC rsALFF | 0.368 | 0.308 | 0.543 | 0.014 | 0.582 | 0.004 | 0.445 | 0.104 | 0.399 | 0.216 | 0.448 | 0.136 |
| L HPC volume (VBM cluster) | 0.360 | 0.308 | 0.449 | 0.072 | 0.495 | 0.024 | 0.351 | 0.432 | 0.401 | 0.190 | 0.467 | 0.096 |
| R thalamus volume (VBM cluster) | 0.361 | 0.308 | 0.422 | 0.110 | 0.373 | 0.253 | 0.249 | > 0.999 | 0.338 | 0.350 | 0.463 | 0.099 |
| R-L HPC rsFC | 0.498 | 0.039 | 0.218 | > 0.999 | 0.332 | 0.352 | 0.313 | 0.781 | 0.122 | > 0.999 | 0.107 | > 0.999 |
| M thalamus volume (VBM cluster) | 0.397 | 0.180 | 0.326 | 0.416 | 0.356 | 0.290 | 0.223 | > 0.999 | 0.179 | 0.933 | 0.463 | 0.099 |
| PrCu rsALFF | 0.328 | 0.464 | 0.425 | 0.140 | 0.374 | 0.290 | 0.171 | > 0.999 | 0.281 | 0.594 | 0.460 | 0.126 |
| R HPC volume (VBM cluster) | 0.276 | 0.549 | 0.355 | 0.297 | 0.334 | 0.352 | 0.243 | > 0.999 | 0.556 | 0.008 | 0.413 | 0.147 |
| R HPC volume (segmentation) | 0.280 | 0.549 | 0.258 | 0.906 | 0.259 | 0.489 | 0.223 | > 0.999 | 0.508 | 0.026 | 0.395 | 0.166 |
| R ERC volume (segmentation) | 0.303 | 0.476 | 0.241 | 0.942 | 0.258 | 0.489 | 0.171 | > 0.999 | 0.365 | 0.272 | 0.025 | > 0.999 |
| L HPC volume (segmentation) | 0.281 | 0.549 | 0.166 | > 0.999 | 0.332 | 0.352 | 0.181 | > 0.999 | 0.412 | 0.171 | 0.393 | 0.166 |
| L thalamus (segmentation) | 0.238 | 0.549 | 0.194 | > 0.999 | 0.330 | 0.352 | 0.096 | > 0.999 | 0.288 | 0.594 | 0.558 | 0.015 |
| R HPC -PCC rsFC | 0.281 | 0.549 | 0.106 | > 0.999 | 0.269 | 0.489 | 0.274 | > 0.999 | 0.055 | > 0.999 | 0.291 | 0.528 |
| R HPC - MPFC rsFC | 0.118 | 0.549 | -0.118 | > 0.999 | 0.078 | 0.663 | 0.074 | > 0.999 | 0.290 | 0.594 | 0.227 | 0.738 |

Table S4.2: score-level correction for multiple testing (n=13), separately for each of the different memory scores (n=6) examined; **bold**: p-corr < 0.05; HPC: hippocampus; MAP: Memory and Amnesia Project; MPFC: medial prefrontal cortex; OPTIMA: Oxford Project To Investigate Memory and Aging; PCC: posterior cingulate cortex; p-corr: p values of bivariate correlations are corrected for multiple testing using the Holm-Bonferroni sequential method of correction for the number of different variables (n = 13) per memory score examined; PrCu: precuneus; rsALFF: amplitude of low frequency fluctuations; rsFC: resting-state functional connectivity; VBM: voxel-based morphometry; z-res: Mean rsALFF and rsFC values are residualized for age and sex across participants; volumes (derived from manual / automated segmentation or from the mean GM volume expressed by VBM clusters) were residualized against age, sex, TIV, and scan source (MAP, OPTIMA) across participants.

#### S5: Relationship of memory scores with volumes of manually delineated HPC portions

Anterograde Memory: Verbal Recognition: We found no behavioral correlations with manually delineated anterior/posterior HPC portions at uncorrected levels (all rs, 0.301 ≥ r ≥ 0.171; all ps, 0.312 ≥ p-unc ≥ 0.070).

Anterograde Memory: Visual Recognition: No significant correlations were observed between visual recognition scores and the volumes of the manually delineated left/right anterior/posterior HPC portions at uncorrected levels (all ps, p-unc ≥ 0.080).

Anterograde Memory: Verbal Recall: A weak correlation was observed with the left anterior HPC at uncorrected levels (r = 0.351, p-unc = 0.033; rest of ps, p-unc ≥ 0.095). Since patients’ rsALFF in the PCC did not correlate significantly with the volume of the manually delineated left anterior HPC (r = 0.333, p = 0.050), we entered these two factors as independent variables in a multiple step-wise linear regression, with verbal recall scores as the dependent variable. The regression terminated in a single step, with rsALFF in the PCC as the only predictor of patients’ performance (R^2^=0.34; β(z)=0.58; F=16.40, p < 0.0005).

Anterograde Memory: Visual Recall: Of the manually delineated HPC portions, we observed no correlations with visual recall memory scores at uncorrected levels (all rhos, rho ≤0.272; all ps, p-unc ≥0.109).

Anterograde Memory: Retention (forgetting): There was no clear evidence for a selective relationship of visual forgetting with anterior vs. posterior portions of the HPC (all rhos, 0.471≥ rho ≥ 0.332; all ps, 0.055 ≥ p-unc ≥ 0.005).

Remote autobiographical memory: Correlations of remote autobiographical memory with volumes of manually delineated portions of the HPC (anterior / posterior right / left HPC) were observed at uncorrected levels (0.429 ≥ r ≥ 0.334; 0.066 ≥ p-unc ≥ 0.016). All four HPC portions correlated volumetrically with the left thalamus (right anterior HPC: r = 0.327, p-unc = 0.045; left anterior HPC: r = 0.427, p-unc = 0.007; right posterior HPC: r = 0.362, p-unc = 0.025; left posterior HPC: r = 0.360, p-unc = 0.026). A series of four partial correlational analyses demonstrated that left thalamic volume correlated with remote autobiographical memory scores over and above the volume of each of those four manually delineated HPC portions (control variable: right anterior HPC: r = 0.489, p = 0.006; left anterior HPC: r = 0.482, p = 0.007; right posterior HPC: r = 0.505; p = 0.004; left posterior HPC: r = 0.509, p = 0.004).

| HPC portion | Verbal Recognition (z) | | Visual Recognition (z) | | Verbal Recall (z) | | Visual Recall (z) | | Visual Forgetting (z) | | Remote Autobiographical (18) | |
| --- | --- | --- | --- | --- | --- | --- | --- | --- | --- | --- | --- | --- |
|  | r | p-unc | r | p-unc | r | p-unc | rho | p-unc | rho | p-unc | r | p-unc |
| R aHPC (z-res) | 0.173 | 0.305 | 0.296 | 0.080 | 0.158 | 0.352 | 0.272 | 0.109 | 0.435 | 0.010 | 0.364 | 0.044 |
| L aHPC (z-res) | 0.294 | 0.077 | 0.237 | 0.164 | 0.351 | 0.033 | 0.265 | 0.118 | 0.332 | 0.055 | 0.390 | 0.030 |
| R pHPC (z-res) | 0.301 | 0.070 | 0.240 | 0.159 | 0.278 | 0.095 | 0.195 | 0.254 | 0.461 | 0.006 | 0.429 | 0.016 |
| L pHPC (z-res) | 0.171 | 0.312 | 0.122 | 0.480 | 0.250 | 0.135 | 0.104 | 0.546 | 0.471 | 0.005 | 0.334 | 0.066 |

Table S5.1: aHPC: anterior hippocampus; HPC: hippocampus; L: left hemisphere; MAP: Memory and Amnesia Project; OPTIMA: Oxford Project To Investigate Memory and Aging; pHPC: posterior hippocampus; p-unc: p values of bivariate correlations are presented at uncorrected levels for display purposes; R: right hemisphere; rho: Spearmann’s rank correlation coefficient; r: Pearson’s correlation coefficient; TIV: total intracranial volume; z: average of age-scaled standardized scores on neuropsychological tests of episodic memory; z-res: volumes for each HPC portion are residualized against age, sex, TIV, and scan source (MAP, OPTIMA) across participants.

#### S6: ‘Impaired’ vs. ‘Unimpaired’ patients on visual forgetting: Structural/Functional abnormalities

Given that 17 of our patients reached ceiling scores in visual forgetting, we also dichotomized our patient group into two subgroups, those that attained ceiling scores (z=0.33), and those with lower scores (z<0.33). We therefore compared the two patient subgroups across the 13 structural and functional abnormalities identified above at whole-group level. Consistent with our correlational approach, the two patient subgroups differed only with respect to the volume of the right HPC, as expressed in manual volumetry (t = -3.32, p-corr = 0.027) and in the right HPC VBM cluster (t = -4.05, p-corr = 0.004; rest of ps, p-corr ≥ 0.143).

| Structural / Functional abnormalities | ‘Impaired’ patients (z < 0.33) | | ‘Unimpaired’ patients | | t/Wt | p-corr | F | p-corr |
| --- | --- | --- | --- | --- | --- | --- | --- | --- |
|  | Mean (z-res) | SD (z-res) | Mean (z-res) | SD (z-res) |  |  |  |  |
| R HPC volume (VBM) | -1.31 | 0.84 | -0.01 | 1.00 | -4.05 | 0.004 | 14.94 | 0.0075 |
| R HPC volume (segmentation) | -1.27 | 0.81 | -0.15 | 1.09 | -3.32 | 0.027 | 9.96 | 0.0480 |
| L HPC volume (VBM) | -1.01 | 1.16 | -0.07 | 1.07 | -2.43 | 0.190 | 4.20 | 0.3053 |
| L HPC volume (segmentation) | -1.15 | 0.96 | -0.17 | 1.16 | -2.63 | 0.143 | 5.60 | 0.2500 |
| R ERC volume (segmentation) | -0.79 | 0.91 | 0.07 | 1.06 | -2.48 | 0.190 | 4.45 | 0.3053 |
| R thalamus volume (VBM) | -0.96 | 1.04 | -0.19 | 0.81 | -2.43 | 0.190 | 5.34 | 0.2520 |
| M thalamus volume (VBM) | -0.76 | 0.74 | -0.38 | 0.90 | -1.33 | 0.645 | 1.38 | > 0.999 |
| L thalamus volume (segmentation) | -0.68 | 0.88 | -0.15 | 1.04 | -1.56 | 0.645 | 1.08 | > 0.999 |
| PCC rsALFF | -0.84 | 0.75 | -0.15 | 0.88 | -2.36 | 0.190 | 6.00 | 0.2319 |
| PrCu rsALFF | -0.83 | 0.75 | -0.21 | 0.89 | -2.10 | 0.270 | 5.36 | 0.2520 |
| R HPC – MPFC rsFC | -0.79 | 0.57 | -0.38 | 0.85 | -1.53 | 0.645 | 1.92 | 0.8871 |
| R HPC - PCC rsFC | -0.62 | 0.59 | -0.37 | 1.07 | -0.84 | 0.816 | < 0.005 | > 0.999 |
| R-L HPC rsFC | -0.58 | 0.92 | -0.38 | 1.17 | -0.51 | 0.816 | 0.19 | > 0.999 |

Table S6.1: ‘Impaired’ subgroup: patients that scored below the ceiling level (z < 0.33) for Visual Forgetting in the Doors and People test (Baddeley et al., 1994); Unimpaired subgroup (n=17): patients that scored at ceiling level (z=0.33) p-corr: significance values are corrected for multiple testing using the Holm-Bonferroni sequential method (Holm, 1979); t/Wt: comparison between the two subgroups across the 13 structural/functional abnormalities identified at group level for patients; F: these comparisons were iterated in the form of a series of univariate ANCOVAs, including patients’ scores for depression (HADS) as between-subjects covariates; z-res: volumes are residualized against age, sex, scan source (MAP, OPTIMA), and TIV; functional abnormalities are residualized against age and sex; t: Student t-test; Wt: Welch t-test; SD: standard deviation; **bold**: p-corr < 0.05; HADS: Hospital Anxiety and Depression Scale (Zigmond and Snaith, 1983); ANCOVA: analysis of covariance; OPTIMA: Oxford Project To Investigate Memory and Aging; Memory and Amnesia Project.

We then iterated these comparisons with a series of one-way ANCOVAs, including patients’ HADS scores for depression as a covariate of no interest. The results retained their significance (manually delineated right HPC volume: F = 9.96; p-corr = 0.048; right HPC VBM cluster: F = 14.94, p-corr = 0.007; rest of ps, p-corr ≥ 0.232).

#### S7: Relationship of Depression with memory impairment and structural / functional abnormalities

Of the three composite memory scores, anterograde retention correlated with scores for depression (HADS) across patients (rho = -0.425, p-corr = 0.045; rest of ps, p-corr > 0.248). We thus also examined the relationship among scores for depression, anterograde retention and HPC atrophy, with which anterograde retention scores strongly correlated. No correlation of scores for depression with HPC volumes (VBM clusters, manually delineated volumes) reached significance even at uncorrected levels (all rhos, |rho| ≤ 0.25; all ps, p-unc ≥ 0.158), and right HPC volumes correlated with visual forgetting scores over and above depression scores (right HPC VBM cluster: rho=0.564, p=0.001; manually delineated right HPC: rho=0.530, p=0.002).

#### S8: Comparison of treated vs. untreated patients on memory scores and structural/functional brain abnormalities

We investigated whether patients that had been treated with immunosuppressive therapy (n=31) showed less pronounced memory impairment and brain abnormalities as compared with those that had not received treatment (n=7), in order to ensure that the structure/function-behavior relationships disclosed across all 38 patients were not driven by the subgroup of patients that had not received such treatment. Patients who had not received such treatment scored lower than the rest of the patients on anterograde retrieval (Wt = 4.49, p-corr = 0.0008), but not on visual forgetting (t = 1.29, p-corr = 0.416), or remote autobiographical memory (t = 1.11, p-corr = 0.416). Nevertheless, the two patient subgroups did not differ with respect to the extent of structural or functional brain abnormalities that we observed above for the entire group of patients (all ps, p-corr ≥ 0.832).

| **Memory Scores** | **Treated (n=31)** | | **Untreated (n=7)** | | **Treated vs Untreated** | |
| --- | --- | --- | --- | --- | --- | --- |
|  | **Mean** | **SD** | **Mean** | **SD** | **t/Wt** | **p-unc** |
| Verbal Recognition (z) | -0.56 | 0.95 | -1.44 | 0.74 | 2.14 | 0.040 |
| Visual Recognition (z) | -0.33 | 1.18 | -1.47 | 1.02 | 2.19 | 0.036 |
| Verbal Recall (z) | -0.90 | 0.75 | -1.65 | 0.29 | 2.39 | 0.022 |
| Visual Recall (z) | -0.61 | 1.36 | -2.29 | 0.60 | 4.81 | < 0.0005 |
| Visual Forgetting (z) | -0.42 | 1.11 | -1.13 | 1.43 | 1.29 | 0.208 |
| Remote Autobiographical Memory (max=18) | 10.19 | 4.74 | 7.60 | 5.13 | 1.11 | 0.278 |
| **Structural/Functional abnormalities (z-res)** | | | | | | |
| R-L HPC rsFC | -0.43 | 1.02 | -0.69 | 1.20 | 0.50 | 0.619 |
| PrCu rsALFF | -0.41 | 0.88 | -0.97 | 0.52 | 1.37 | 0.179 |
| PCC resALFF | -0.38 | 0.88 | -0.86 | 0.53 | 1.16 | 0.255 |
| R HPC - MPFC rsFC | -0.58 | 0.78 | -0.76 | 0.68 | 0.49 | 0.627 |
| R HPC - PCC rsFC | -0.50 | 0.90 | -0.64 | 0.62 | 0.32 | 0.749 |
| R ERC volume (segmentation) | -0.28 | 1.07 | -0.43 | 0.96 | 0.34 | 0.740 |
| L HPC volume (VBM) | -0.42 | 1.14 | -1.34 | 1.31 | 1.89 | 0.067 |
| R Thalamus volume (VBM) | -0.55 | 1.01 | -0.57 | 1.09 | 0.05 | 0.958 |
| R HPC volume (VBM) | -0.50 | 1.12 | -1.35 | 0.90 | 1.86 | 0.071 |
| M Thalamus volume (VBM) | -0.56 | 0.87 | -0.60 | 0.69 | 0.12 | 0.907 |
| L Thalamus volume (segmentation) | -0.37 | 1.03 | -0.53 | 0.58 | 0.38 | 0.703 |
| R HPC volume (segmentation) | -0.53 | 1.10 | -1.36 | 0.70 | 1.91 | 0.064 |
| L HPC volume (segmentation) | -0.47 | 1.16 | -1.26 | 0.90 | 1.69 | 0.100 |

Table S8.1: Patients that had not been treated with immunosuppressive therapy (n=7) were more impaired in tests of anterograde retrieval (visual/verbal recall/recognition) as compared with those that had received immunosuppressive therapy (n=31). The two groups did not differ with respect to structural or functional brain abnormalities, even at uncorrected levels (p-unc); z-res: volumes are residualized against age, sex, scan source (MAP, OPTIMA), and TIV; functional abnormalities are residualized against age and sex.

Given the more pronounced impairment on anterograde retrieval of patients that had not received immunosuppressive therapy, we sought to determine whether the relationship of this composite score with reduced rsALFF in the PCC held when the analysis was confined to the 31 patients that had received immunosuppressive therapy. Indeed, the relationship between reduced rsALFF in the PCC and impaired anterograde retrieval retained its significance across this patient subgroup (r = 0.532, p-corr = 0.032; rest of ps, p-corr ≥ 0.228).

| **Structural/Functional Abnormalities (z-res)** | **r** | **p-corr** |
| --- | --- | --- |
| PCC rsALFF | 0.538 | 0.028 |
| L HPC volume (VBM) | 0.427 | 0.204 |
| R thalamus volume (VBM) | 0.401 | 0.275 |
| R-L HPC rsFC | 0.390 | 0.330 |
| M thalamus volume (VBM) | 0.362 | 0.405 |
| R ERC volume (segmentation) | 0.292 | 0.888 |
| PrCu rsALFF | 0.274 | 0.888 |
| R HPC volume (VBM) | 0.258 | 0.888 |
| R HPC - PCC rsFC | 0.287 | 0.888 |
| L Thalamus volume (segmentation) | 0.217 | 0.964 |
| R HPC volume (segmentation) | 0.160 | > 0.999 |
| L HPC volume (segmentation) | 0.165 | > 0.999 |
| R HPC - MPFC rsFC | -0.007 | > 0.999 |

Table S8.2: Anterograde retrieval composite scores (mean of age-scaled standardized scores on tests of visual / verbal recall/recognition) correlated with the mean rsALFF in the PCC across the 31 patients that had received immunosuppressive therapy; L, R: left, right (hemisphere); HPC: hippocampus; MPFC: medial prefrontal cortex; p-corr: p values are adjusted for multiple testing using the Holm-Bonferroni sequential method; **bold**: p-corr < 0.05; rsALFF: resting-state amplitude of low frequency fluctuations; rsFC: resting-state functional connectivity; PrCu: precuneus; PCC: posterior cingulate cortex; z-res: volumes are residualized against age, sex, scan source (MAP, OPTIMA), and TIV; functional abnormalities are residualized against age and sex;

#### S9: Participants: Data availability

| **Group** | **Project** | **N** | **Structural MRI** | **Resting-state fMRI** | **Neuropsychological assessment** | **Comments** |
| --- | --- | --- | --- | --- | --- | --- |
| Healthy  Controls | OPTIMA | 32 | ✔ | × | × | No rsfMRI or neuropsychological data* available |
|  | MAP | 28 | ✔ | ✔ | ✔ | Full protocol |
|  |  | 1 | ✔ | × | ✔ | Full protocol – rsfMRI datasets discarded due to acquisition errors and/or movement |
|  |  | 4 | ✔ | ✔ | × | No neuropsychological assessment due to scheduling conflicts |
|  |  | 2 | ✔ | × | × | No neuropsychological assessment due to scheduling conflicts- rsfMRI datasets discarded due to acquisition errors and/or movement |
|  |  | 12 | × | × | ✔ | No structural / functional MRI due to scheduling conflicts |
| Patients |  | 35 | ✔ | ✔ | ✔ | Full protocol |
|  |  | 3 | ✔ | × | ✔ | Full protocol – rsfMRI datasets discarded due to acquisition errors and/or movement |

Table S9.1: Numbers of healthy controls and patients that underwent structural, functional MRI, and neuropsychological assessment. *: While the healthy controls whose structural MRI datasets were added from the OPTIMA project had not been assessed with our laboratory’s neuropsychological battery, they had been assessed with tests measuring overall cognitive impairment [Mini-Mental State Examination – MMSE; (Folstein et al., 1975)]. Expectedly, these scores indicated that none of those healthy controls had any apparent cognitive impairment [mean = 29.74, SD = 0.56, min = 28, well above widely accepted cut-offs, e.g. (Aevarsson and Skoog, 2000; Di Carlo et al., 2002)]. OPTIMA: Oxford Project To Investigate Memory and Aging; Memory and Amnesia Project; (rs)fMRI: (resting-state functional) Magnetic Resonance Imaging; n: number of participants.
